# Downregulating PTBP1 fails to convert astrocytes into hippocampal neurons and to alleviate symptoms in Alzheimer’s mouse models

**DOI:** 10.1101/2022.04.27.489696

**Authors:** Tiantian Guo, Xinjia Pan, Guangtong Jiang, Denghong Zhang, Jinghui Qi, Lin Shao, Zhanxiang Wang, Huaxi Xu, Yingjun Zhao

## Abstract

Conversion of astroglia into functional neurons has been considered as a promising therapeutic strategy for neurodegenerative diseases. Recent studies reported that downregulation of the RNA binding protein, PTBP1, converts astrocytes into neurons in situ in multiple mouse brain regions, consequently improving pathological phenotypes associated with Parkinson’s disease, RGC loss, and aging. Here, we demonstrate that PTBP1 downregulation using antisense oligonucleotides or an astrocyte specific AAV-mediated shRNA system fails to convert hippocampal astrocytes into neurons in WT, and β-amyloid (5×FAD) and tau (PS19) Alzheimer’s disease (AD) mouse models, and fails to reverse synaptic/cognitive deficits and AD-associated pathology. Similarly, PTBP1 downregulation cannot convert astrocytes into neurons in the striatum and substantia nigra. Together, our study suggests that cell fate conversion strategy for neurodegenerative disease therapy through manipulating one single gene, such as PTBP1, warrants more rigorous scrutiny.

## Introduction

Neuronal loss is the primary cause for functional deterioration in the central nervous system (CNS) in neurodegenerative diseases such as Parkinson’s disease (PD) and Alzheimer’s disease (AD) (Kalia & Lang, 2015; Mesulam, 1999). Replenishment of neurons is considered as an effective approach to restore CNS function. Two major regeneration strategies utilizing neuronal lineage cells have been developed so far: transplantation of exogenous iPSC-derived neuronal cells and activation of endogenous neurogenesis niche. However, transplantation of exogenous cells may induce immunological rejection and tumorigenesis (Olanow et al., 2003; Trounson & McDonald, 2015). Adult neurogenesis only occurs in a few small brain regions in rodent models, and whether it exists in humans remains controversial (Gage, 2019; Kempermann et al., 2018). In pursuing novel approaches for neuronal replenishment, attempts have been made to transdifferentiate residential glia cells into functional neurons (Chen et al., 2020; Grande et al., 2013; Guo et al., 2014; Lentini et al., 2021; Liu et al., 2015; Maimon et al., 2021; Matsuda et al., 2019; Mattugini et al., 2019; Niu et al., 2015; Qian et al., 2020; Tai et al., 2021; Torper et al., 2015; Zhou et al., 2020). The *in situ* glia-to-neuron conversion represents an ideal strategy for neuronal regeneration, because: (1) glial cells including astrocytes and microglia are over-proliferated and become reactive (this process was termed as gliosis), leading to neuroinflammation during neurodegeneration; (2) neurons can be regenerated from the excessive glia in designated brain regions where neuronal loss occurs.

Among the targets for converting astroglia into neurons, polypyrimidine tract-binding protein 1 (PTBP1) has attracted much attention, because downregulation of this single protein has been indicated to be sufficient to directly convert astrocytes into functional neurons with high efficiencies, within several weeks to months (Maimon et al., 2021; Qian et al., 2020; Zhou et al., 2020). Specifically, downregulation of PTBP1 by AAV-mediated shRNA system or antisense oligonucleotides (ASO) converted midbrain astrocytes to dopaminergic neurons, and reversed motor deficits in a chemically induced PD mouse model (Qian et al., 2020). Similar motor function improvement was observed in the PD mice when striatal astrocytes were converted into functional neurons by CRISPR-CasRx-mediated PTBP1 downregulation (Zhou et al., 2020). In addition, injection of ASO targeting *Ptbp1* into cerebral spinal fluid (CSF) generated new functional cortical or hippocampal neurons in both young and aged mice (Maimon et al., 2021). Those results, should they sustain, would represent revolutionary advancement in therapeutics for neurodegenerative diseases.

In the current study, utilizing ASO and an astrocyte-restrictive AAV shRNA system, we examine whether the reported astrocyte-to-neuron conversion induced by PTBP1 downregulation can happen in the hippocampus and consequently rescue synaptic and cognitive impairments in AD-associated mouse models, namely amyloid and tau (neurofibrillary tangle) models, 5×FAD and PS19, respectively. In addition, we re-examine whether PTPB1 knockdown can efficiently convert astrocytes into neurons in the brain regions of substantia nigra and striatum, as previously reported.

## Results

### Downregulation of PTBP1 by ASO fails to promote the generation of neurons in mouse hippocampus

Because there is a great potential for ASO in clinical applications (Leavitt & Tabrizi, 2020; Miller et al., 2020; Miller et al., 2013), we first tested the effect of *Ptbp1* ASO (with the same nucleotide sequence as reported in (Qian et al., 2020) on neuronal generation. Two weeks after mouse hippocampal injection of a FAM-labeled ASO targeting *Ptbp1 (Ptbp1-ASO),* we observed FAM signals in mostly NeuN^+^ and to a much lesser extent GFAP^+^ cells near the injection site (**Figure 1—figure supplement 1A**), indicating the lack of astrocytic specificity for ASO delivery. *Ptbp1-ASO* injection resulted in a significant reduction, but incomplete elimination of PTBP1 immunofluorescent (IF) intensities in GFAP^+^cells (**Figure 1—figure supplement 1**, **B** and **C**); however, the number of cells labeled with doublecortin X (DCX), a marker for neural progenitor cells and immature neurons, and the area labeled by mature neuronal marker NeuN are indistinguishable between the *Ptbp1-ASO* and the control groups (**Figure 1— figure supplement 1, D** to **F**). These data suggest that downregulation of PTBP1 by ASO cannot promote the generation of both immature and mature neurons.

### Knockdown of PTBP1 by an astrocyte-restrictive AAV shRNA fails to convert hippocampal astrocytes into neurons *in vivo*

AAV driven by specific promoters has been widely used to efficiently deliver genes or DNA fragments including shRNAs into particular types of CNS cells. We constructed an *hGFAP* promoter-driven AAV vector, which comprises a mCherry reporter and a *Ptbp1* -targeting shRNA in a miR30 cassette (**Figure 1A**). A modified longer form of *hGFAP* promoter was used to ensure the specificity of astrocytic expression, as many serotypes of the AAV driven by the short form *hGFAP* promoter have been reported to having leaky expression in non-astrocytic cells (Wang et al., 2021). The knockdown efficiency of three AAV shRNA viruses was compared with the control AAV scramble virus in primary astrocytic cultures. Because of the highest knockdown efficiency of AAV-GFAP-sh*Ptbp1*-1 (**Figure 1**, **B** and **C**) (nucleotide sequence of the shRNA is identical to that used in (Qian et al., 2020)), it was chosen and designated as AAV-*shPtbp1*or *shPtbp1* for all subsequent experiments.

**Figure 1.**
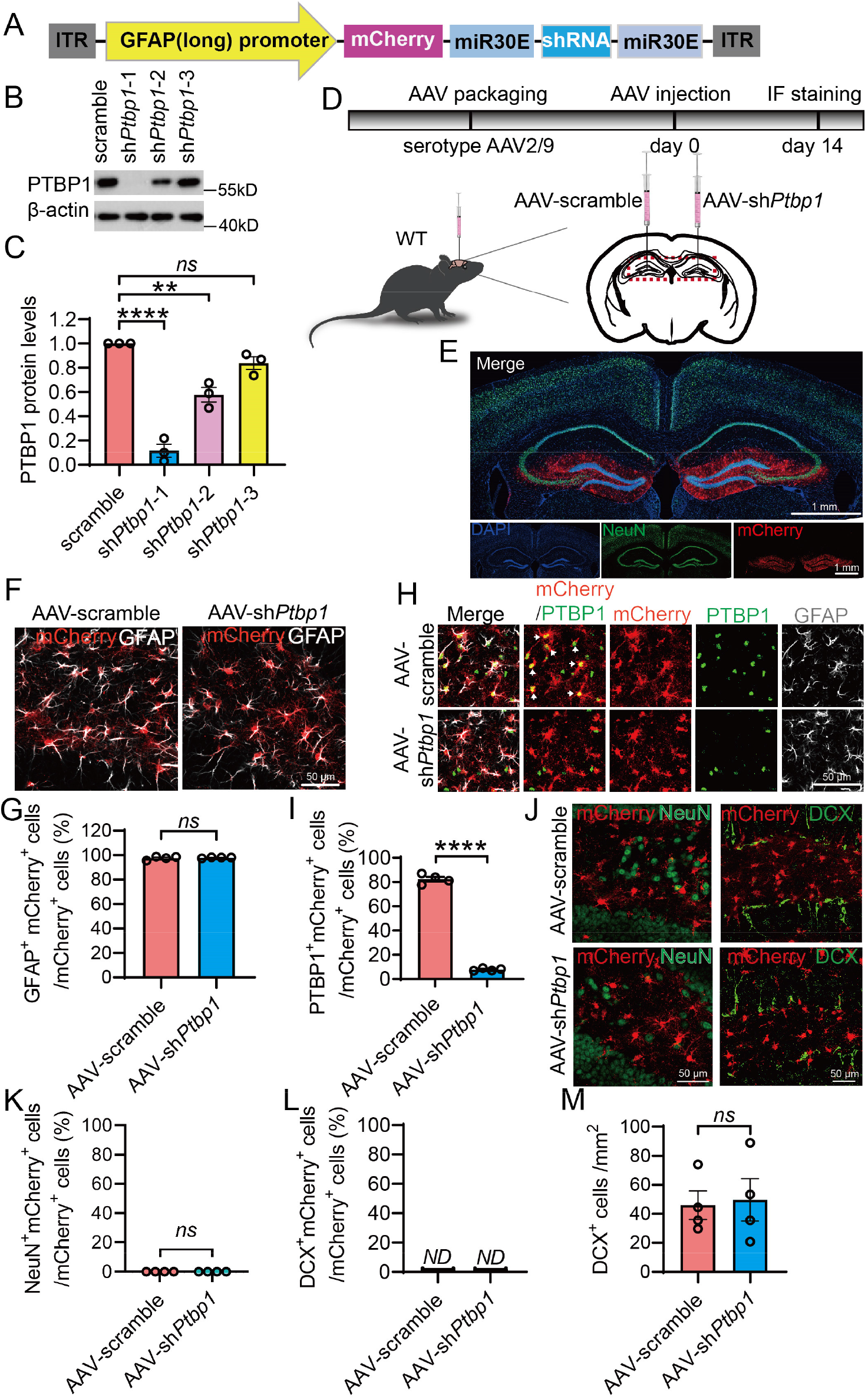
Specific knockdown of PTBP1 in astrocytes fails to convert hippocampal astrocytes into neurons in WT mice. (**A**) Schematic of the AAV-shRNA vector used in this study. (**B** and **C**) Western blot analysis of PTBP1 expression in mouse primary astrocytes 7 days post AAV transduction. (B) Representative PTBP1 blots, (C) quantification. n = 3 independent experiments, one-way ANOVA with Tukey’s multiple comparisons. (**D**) Flow chart of study design to assess PTBP1 downregulation in mouse hippocampus. (**E**) Confocal images of the mouse hippocampus and cortex 2 weeks after the AAV injection. (**F** and **G**) Confocal analysis of mCherry expression in GFAP^+^ cells. (F) Representative images, (G) quantification. n = 4 animals, unpaired t-test. (**H** and **I**) Confocal analysis of PTBP1 expression in mCherry^+^ cells. (H) Representative images, (I) quantification. n = 4 animals, unpaired t-test. (**J** to **M**) Confocal analysis of fluorescent cells. (J) Representative images, (K-M) quantification. n = 4 animals, unpaired t-test. All quantified data are represented as mean ± SEM; **p <0.01; ****p <0.0001; *ns*, not significant; *ND,* undetectable. Figure 1— source data 1 to 7, source data for Figure 1B, 1C, 1G, 1l, 1K, 1L, 1M.

We next transduced AAV-sh *Ptbp1* unilaterally into the right hippocampal dentate gyrus, and AAV-scramble control in the left dentate gyrus, of adult WT mice (**Figure 1D**). Two weeks after the transduction, both sides showed restricted mCherry expression in GFAP^+^ astrocytes in the dentate gyrus and the nearby areas (**Figure 1, E** to **G**). IF staining for PTBP1 showed that AAV-*shPtbp1* achieved nearly complete depletion of PTBP1 in mCherry^+^ cells, and consequently dramatic reduction in the number of PTBP1^+^mCherry^+^ cells, when compared to the control side (**Figure 1, H** and **I**). These results demonstrate that AAV-sh*Ptbp1* can rapidly, efficiently, and specifically knockdown PTBP1 in the hippocampal astrocytes *in vivo.* Nevertheless, AAV-sh*Ptbp1* yielded only handful and neglectable numbers of NeuN^+^mCherry^+^ cells, which are similar to AAV-scramble control (**Figure 1, J** and **K**). DCX^+^mCherry^+^ cells were undetectable, whereas the numbers of DCX^+^ cells were comparable in the knockdown and control groups (**Figure 1, J, L** and **M**). Our data clearly show that suppression of PTBP1 expression cannot convert astrocytes into either immature or mature neurons.

### Sustained downregulation of PTBP1 fails to convert hippocampal astrocytes into neurons in an amyloid AD mouse model

Next, we investigated whether a long-term and continuous PTBP1 downregulation can enhance neuronal generation in 5×FAD mice, a widely used AD mouse model with progressive amyloid-β (Aβ) pathology, synaptic and cognitive impairments (Guo et al., 2020). The long-term knockdown of PTBP1 strategy was based on the fact that the astrocyte-to-neuron conversion as well as the pathogenesis of most neurodegenerative diseases such as AD and PD are progressive and chronic. We injected AAV-sh*Ptbp1* or AAV-scramble bilaterally into the hippocampus of 5×FAD and WT control mice at ~5.5 months of age, and performed IF analyzes at 1 and up to 3 months after the transduction. AAV-sh*Ptbp1* markedly reduced the expression of PTBP1 in mCherry^+^ cells, when compared to the AAV-scramble group (**Figure 2, A** and **B,** and **Figure 2—figure supplement 1, A** and **B**). However, only a neglectable fraction of NeuN^+^mCherry^+^ cells were observed, whereas the vast majority of mCherry^+^ cells are GFAP^+^ astrocytes (**Figure 2, C** to E, and **Figure 2—figure supplement 1, C** to **E**). In addition, NeuN^+^ areas did not vary between AAV-*shPtbp1* and AAV-scramble groups (**Figure 2, C** and **F,** and **Figure 2—figure supplement 1, C** and **F**). Further, AAV-sh*Ptbp1* failed to reverse the over-proliferation of GFAP^+^ astrocytes and the reduction of DCX^+^ cells in 5×FAD mice (**Figure 2, C** and **G** to **I,** and **Figure 2—figure supplement 1, C** and **G to I**), two phenomena that have been reported (Demars et al., 2010; Perez-Nievas & Serrano-Pozo, 2018; Zaletel et al., 2018). DCX^+^mCherry^+^ cells were completely undetectable in all groups (**Figure 2, H** and **J** and **Figure 2—figure supplement 1, H** and **J**). Together, these results demonstrate that sustained downregulation of PTBP1 in the hippocampus also fails to convert astrocytes into neurons in 5×FAD mice.

**Figure 2.**
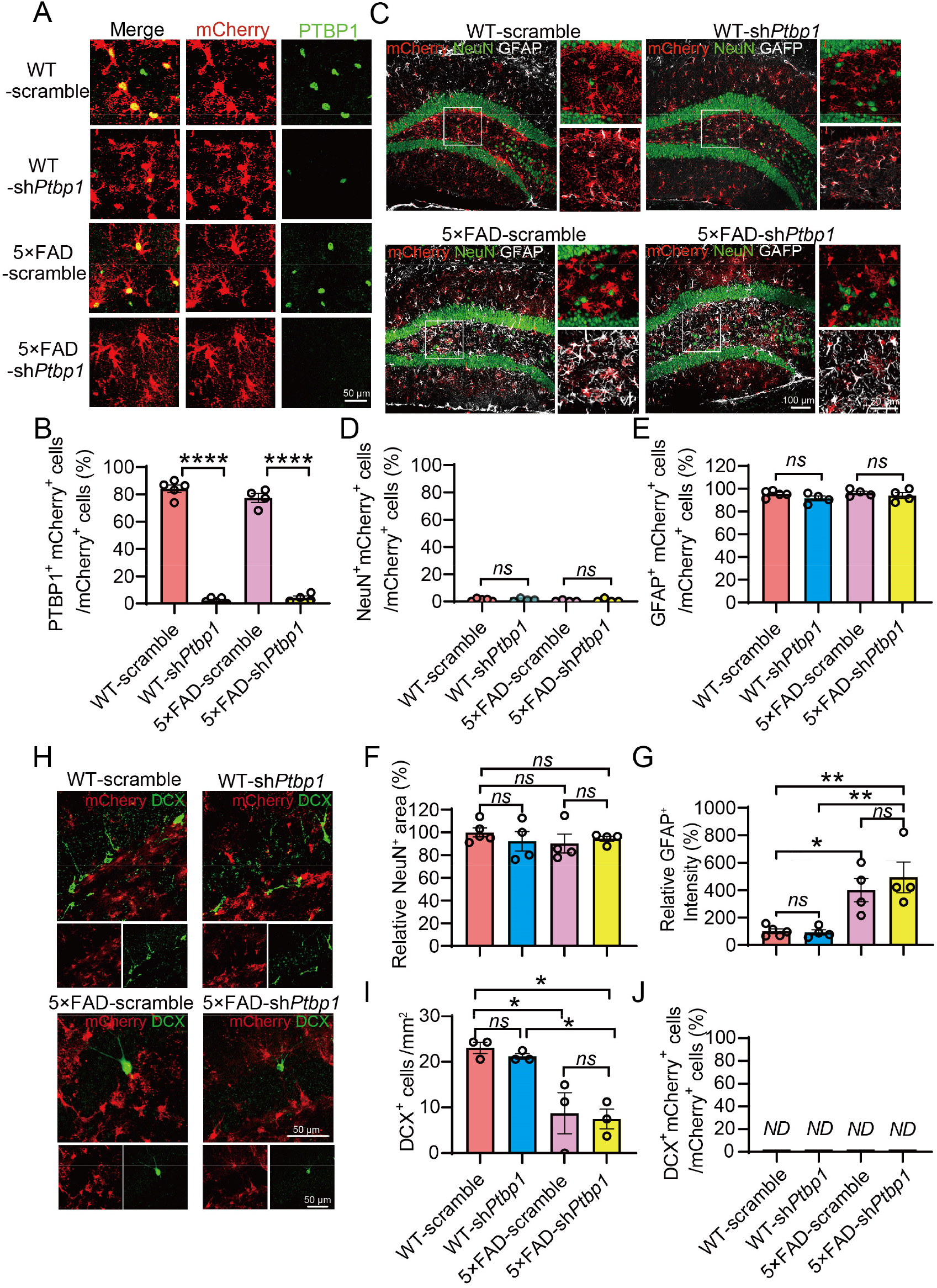
Downregulation of PTBP1 is not able to convert astrocytes into neurons in the hippocampus of 5×FAD mice. (**A** and **B**) Confocal analysis of PTBP1 expression in mCherry^+^ cells in the hippocampus of 5×FAD and WT control mice 3 months after *shPtbp1* or the scramble control AAV transduction. (A) Representative images, (B) quantification. n = 4 to 5 animals, unpaired t-test. (**C** to **G**) Confocal analysis of fluorescent cells. (C) Representative images, (D-G) quantification. n = 4 to 5 animals, one-way ANOVA with Tukey’s multiple comparisons. (**H** to **J**) Confocal analysis of mCherry^+^ cells and DCX^+^ cells. (H) Representative images (I and J) quantification. n = 3 animals per group, one-way ANOVA with Tukey’s multiple comparisons. All quantified data are represented as mean ± SEM; *p <0.05; **p <0.01; ****p <0.0001; *ns,* not significant; *ND,* undetectable. Figure 2—source data 1 to 7, source data for Figure 2B, 2D, 2E, 2F, 2G, 2I, 2J.

### PTBP1 downregulation cannot alleviate Aβ-associated synaptic and cognitive deficits and pathologies in AD mice

The ultimate goal for neuronal regeneration is to restore brain functions which are impaired in neurodegeneration. We performed behavioral tests to evaluate cognitive functions of the mice after AAV-mediated PTBP1 knockdown. In the Morris water maze test, AAV-sh*Ptbp1* failed to alleviate the deficits of spatial memory in 5×FAD mice in the training and probe test phases (**Figure 3, A** and **B**). Similarly, the novel object recognition (NOR) test showed that comparing to the WT control mice, 5×FAD mice spent less time on the novel object, and AAV-*shPtbp1* was unable to reverse this deficit (**Figure 3C**). As expected, AAV-*shPtbp1* also failed to rescue impairments in long-term potentiation (LTP) and synaptophysin (SYP) /PSD95 labeled synaptic clusters in 5×FAD mice (**Figure 3, D** to **G**). Additionally, amyloid deposition in the hippocampus of 5×FAD mice was not altered after PTBP1 downregulation (**Figure 3, H** and **I**). Taken together, PTBP1 downregulation can neither restore synaptic and cognitive function, nor reduce amyloid pathology in 5×FAD mice.

**Figure 3.**
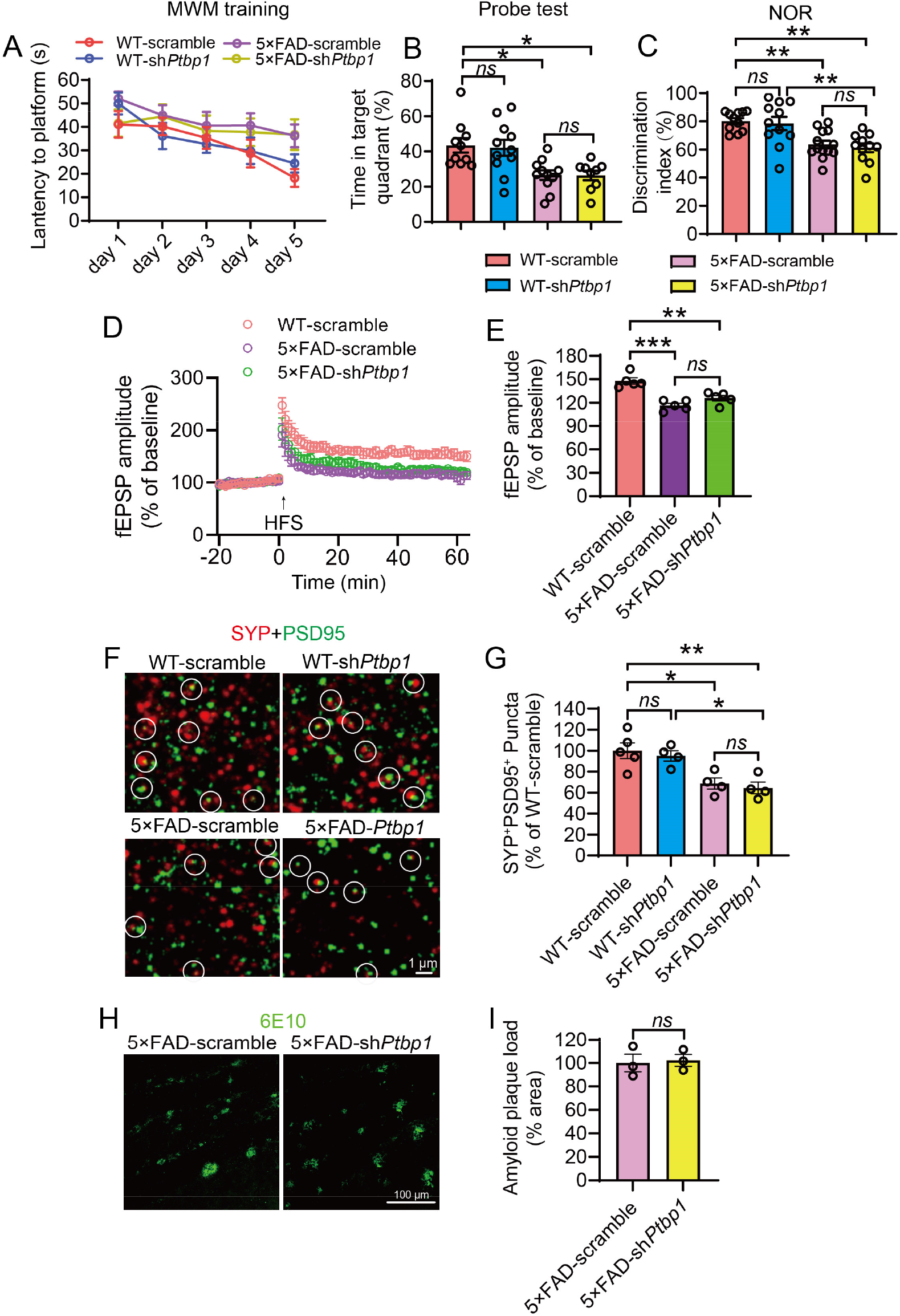
Downregulation of PTBP1 in the hippocampus fails to attenuate cognitive and synaptic deficits as well as Aβ deposition in 5×FAD mice. (**A** and **B**) MWM analysis of 5×FAD and WT control mice 2.5 months after sh*Ptbp1* or the scramble control AAV transduction. (A) The Latency to find the hidden platform during the training phase, (B) time spent in the target quadrant in the probe test. n = 9 to 11 animals, one-way ANOVA with Tukey’s multiple comparisons. (**C**) Quantification of discrimination indexes in the NOR test of the experimental mouse. n = 11 to 13 animals, one-way ANOVA with Tukey’s multiple comparisons. (**D** and **E**) Electrophysiological analysis of LTP. (D) Recordings of hippocampal LTP induced by high-frequency stimulation (HFS), (E) quantification of the field excitatory post synaptic potentials (fEPSP) during the last 10 min of LTP recording. n = 5 brain slices from 4 animals per group, one-way ANOVA with Tukey’s multiple comparisons. (**F** and **G**) Confocal analysis of SYP^+^PSD95^+^ synaptic puncta. (F) Representative images, (G) quantification. n = 4 to 5 animals, one-way ANOVA with Tukey’s multiple comparisons. (**H** and **I**) Confocal analysis of 6E10 (Aβ antibody) stained amyloid deposits. (H) Representative images, (I) quantification. n = 3 animals per group, unpaired t-test. All quantified data are represented as mean ± SEM; *p <0.05; **p <0.01; ***p <0.001 *ns*, not significant. Figure 3—source data 1 to 7, source data for Figure 3A, 3B, 3C, 3D, 3E, 3G, 3I.

### Knockdown of PTBP1 fails to induce the hippocampal astrocyte-to-neuron conversion and to improve cognitive function in tau transgenic mice

Tau pathology has been linked to the degree of dementia and considered to play a causal role in neuronal loss in tauopathies including AD (Fu et al., 2017; Guo et al., 2020). We therefore assessed the PTPB1 knockdown strategy in tau transgenic PS19 mice that develop neuronal loss and brain atrophy starting from 9 month (Yoshiyama et al., 2007). AAV-sh*Ptbp1* or AAV-scramble viruses were injected bilaterally into the hippocampus of 8-month-old PS19 and WT mice. Similar to the 5×FAD mice tested above, the expression of PTBP1 was dramatically reduced in mCherry^+^ cells 3 months post AAV-sh*Ptbp1* transduction (**Figure 4, A** and **B**). mCherry fluorescent signals were detected mostly in GFAP^+^ astrocytes, sparsely in NeuN^+^ neurons, and were undetectable in DCX^+^ cells, in both AAV-sh*Ptbp1* and AAV-scramble groups (**Figure 4, C** to **E, H** and **I**). Consistent with the previous reports (Yoshiyama et al., 2007), we expectedly observed the apparent loss of NeuN^+^ mature neurons and DCX^+^neuronal progenitors, as well as increased GFAP^+^ astrocytes in PS19 mice, all of which, however, remained completely unaffected following the efficient PTBP1 downregulation (**Figure 4, C, F, G, H** and **J**).

**Figure 4:**
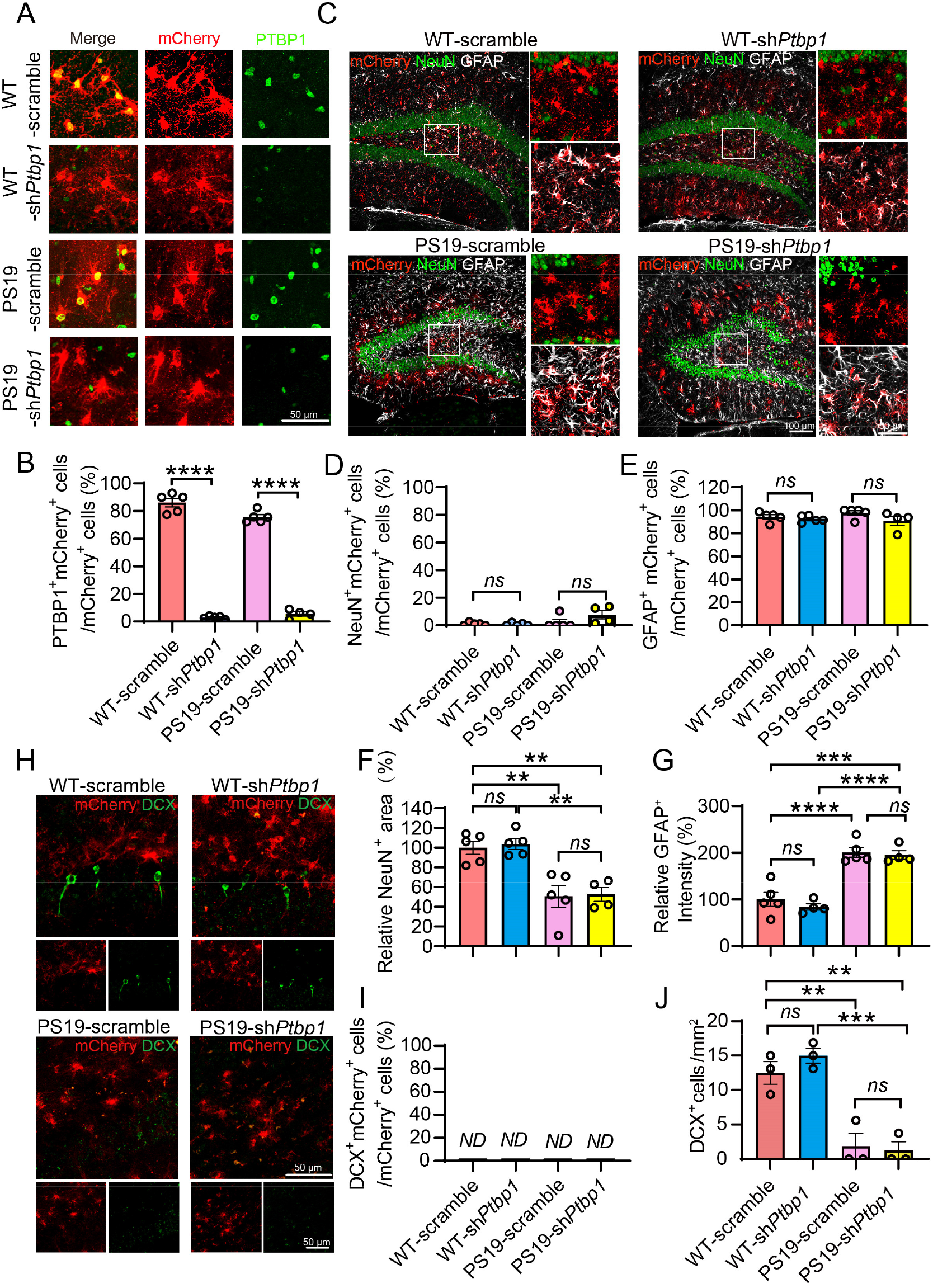
Knockdown of PTBP1 cannot convert astrocytes into neurons in the hippocampus of PS19 mice. (**A** and **B**) Confocal analysis of PTBP1 expression in mCherry^+^ cells in the hippocampus of PS19 and WT control mice 3 months after *shPtbp1* or the scramble control AAV transduction. (A) Representative images, (B) quantification. n = 4 to 5 animals, unpaired t-test. (**C** to **G**) Confocal analysis of fluorescent cells. (C) Representative images, (D-G) quantification. n = 4 to 5 animals, one-way ANOVA with Tukey’s multiple comparisons. (**H** to **J**) Confocal analysis of mCherry^+^ cells and DCX^+^ cells. (H) Representative images, (I and J) quantification. n = 3 animals per group, one-way ANOVA with Tukey’s multiple comparisons. All quantified data are represented as mean ± SEM; **p <0.01; ***p <0.001; ****p <0.0001; *ns,* not significant; *ND,* undetectable. Figure 4—source data 1 to 7, source data for Figure 4B, 4D, 4E, 4F, 4G, 4I, 4J.

Cognitive function of PS19 and WT mice transduced with AAV-sh *Ptbp1* or AAV-scramble were also assessed. In the NOR test, PS19 mice spent significantly less time on the novel object, when compared to the WT control groups; and downregulation of PTBP1 did not improve this performance (**Figure 5A**). We next employed contextual fear conditioning (CFC) test to examine memory decay and found that AAV-sh*Ptbp1* failed to rescue the deficits of PS19 mice in freezing behaviors, when examined two weeks after the initial electric shock (**Figure 5B**). In addition, we surveyed synapses and tau pathology in PS19 mice, and found that loss of SYP^+^PSD95^+^ synaptic clusters and the presence of AT8^+^ phosphor-tau deposition remained and were not alleviated by PTBP1 knockdown (**Figure 5, C** to **F**). The levels of tau phosphorylated at T181 site were also not altered (data not shown).

**Figure 5.**
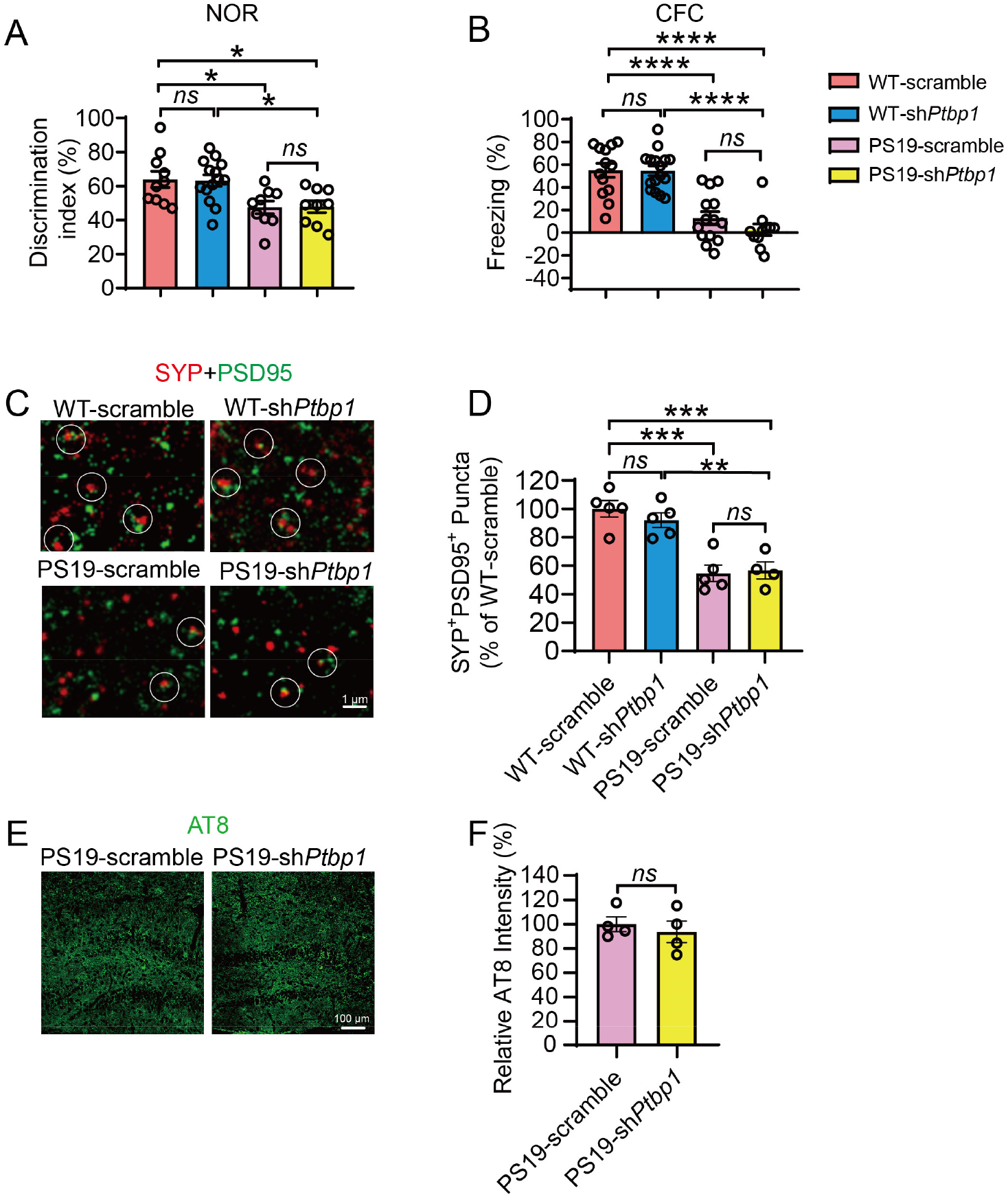
Downregulation of PTBP1 in the hippocampus fails to attenuate cognitive and synaptic deficits as well as tau pathology in PS19 mice. (**A**) Quantification of discrimination indexes in the NOR test of PS19 and WT control mice 2.5 month after the sh*Ptbp1* or the scramble control AAV transduction. n = 9 to 14 animals, one-way ANOVA with Tukey’s multiple comparisons. (**B**) Analysis of the freezing behavior of the experimental mouse during the CFC test. n= 11 to 16 animals, one-way ANOVA with Tukey’s multiple comparisons. (**C** and **D**) Confocal analysis of SYP^+^PSD95^+^ synaptic puncta. (C) Representative images, (D) quantification. n = 4 to 5 animals, one-way ANOVA with Tukey’s multiple comparisons. (**E** and **F**) Confocal analysis of AT8 (phospho-tau antibody) stained tau pathology. (E) Representative images, (F) quantification. n= 4 animals per group, unpaired t-test. All quantified data are represented as mean ± SEM; *p <0.05; **p <0.01; ***p <0.001; ****p <0.0001; *ns*, not significant. Figure 5—source data 1 to 4, source data for Figure 5A, 5B, 5D, 5F.

Together, our results clearly show that downregulation of PTBP1 in hippocampal astrocytes fails to convert astrocytes into neurons, and consequently are unable to improve cognitive function in mice under either physiological or pathological conditions associated with AD.

### Downregulation of PTBP1 fails to convert astrocytes into neurons in either striatum or substantia nigra of mice

To investigate whether the brain-region specificity exists for the presumed astroglia-to-neuron conversion induced by PTBP1 downregulation, we injected AAV-sh*Ptbp1* or AAV-scramble viruses into the striatum or substantia nigra of the adult mice, the two brain regions in which a successful astrocyte-to-neuron conversion by downregulating PTBP1 has been claimed (Qian et al., 2020; Zhou et al., 2020). In our study, similar to the hippocampus, astrocytic PTBP1 expression was nearly completely downregulated by AAV-sh*Ptbp1* in the striatum 1 or 2 months after the viral injection (**Figure 6, A** to **C**). However, we found that only a neglectable portion of mCherry^+^ cells were NeuN^+^, while most of mCherry^+^ cells were GFAP^+^ (**Figure 6, D** to **F**), indicative of a complete failure of astrocyte-to-neuron conversion. In addition, the numbers of NeuN^+^ or GFAP^+^cells near the injection site were indistinguishable between AAV-sh*Ptbp1* and the control AAV-scramble groups (**Figure 6, G** to **H**).

**Figure 6:**
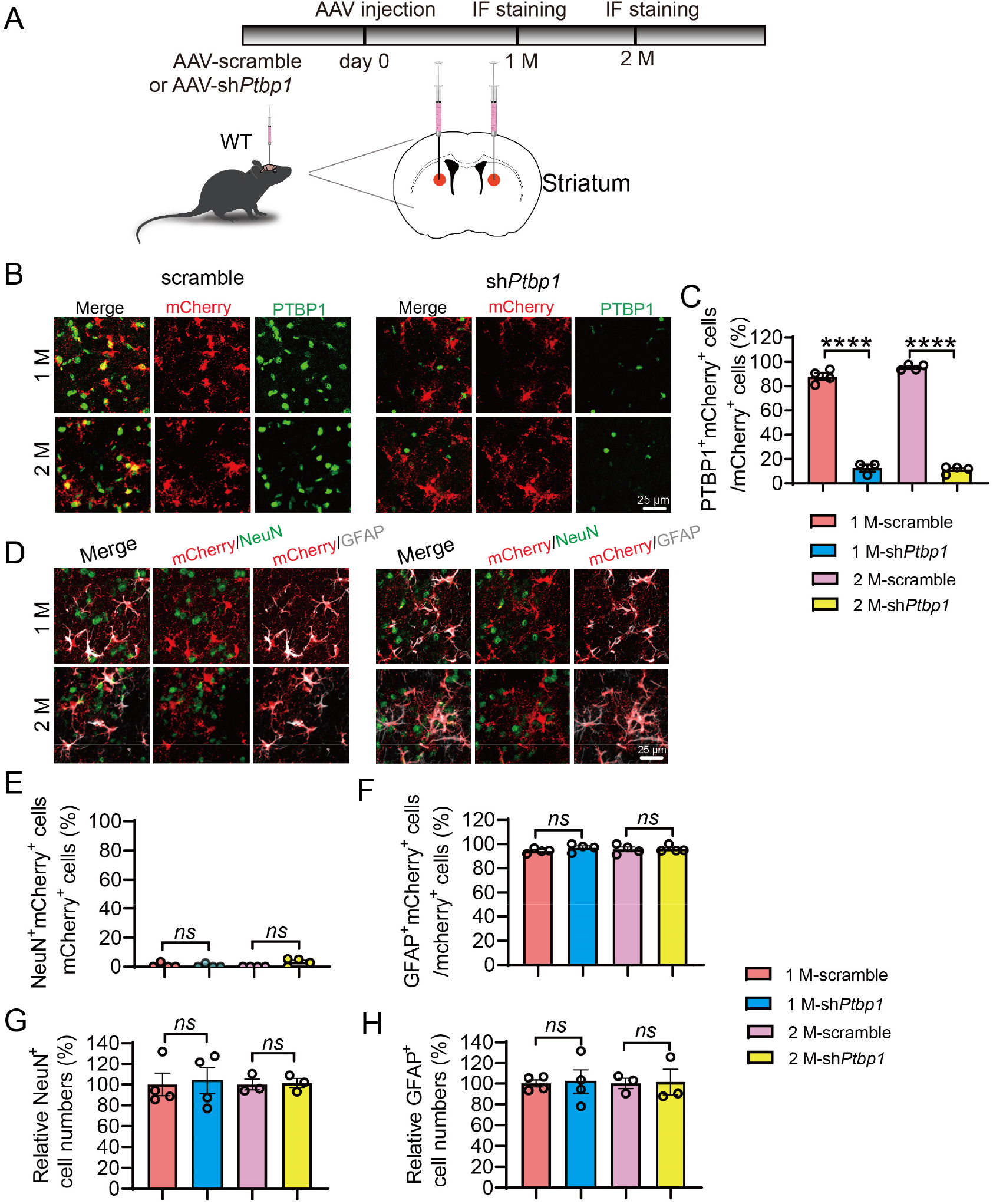
Knockdown of PTBP1 cannot convert astrocytes into neurons in the mouse striatum. (**A**) Schematic of study design. (**B** and **C**) Confocal analysis of PTBP1 expression in mCherry^+^ cells. (B) Representative images, (C) quantification. n = 3 to 4 animals, one-way ANOVA with Tukey’s multiple comparisons. (**D** to **H**) Confocal analysis of fluorescent cells. (D) Representative images, (E-H) Quantification. n = 3 to 4 animals, one-way ANOVA with Tukey’s multiple comparisons. All quantified data are represented as mean ± SEM; ****p <0.0001; *ns*, not significant. Figure 6—source data 1 to 5, source data for Figure 5C, 5E, 5F, 5G, 5H.

Similar results were obtained in the substantia nigra region: an efficient knockdown of PTBP1 in astrocytes did not result in any significant differences in the numbers of NeuN^+^mCherry^+^, GFAP^+^mCherry^+^, NeuN^+^ and GFAP^+^ cells, when compared to the scramble control group (**Figure 7, A** to **H**). Together, our data clearly demonstrate that downregulation of PTBP1 is unable to convert astrocytes into neurons in all brain regions tested including the hippocampus, the striatum and the substantia nigra.

**Figure 7.**
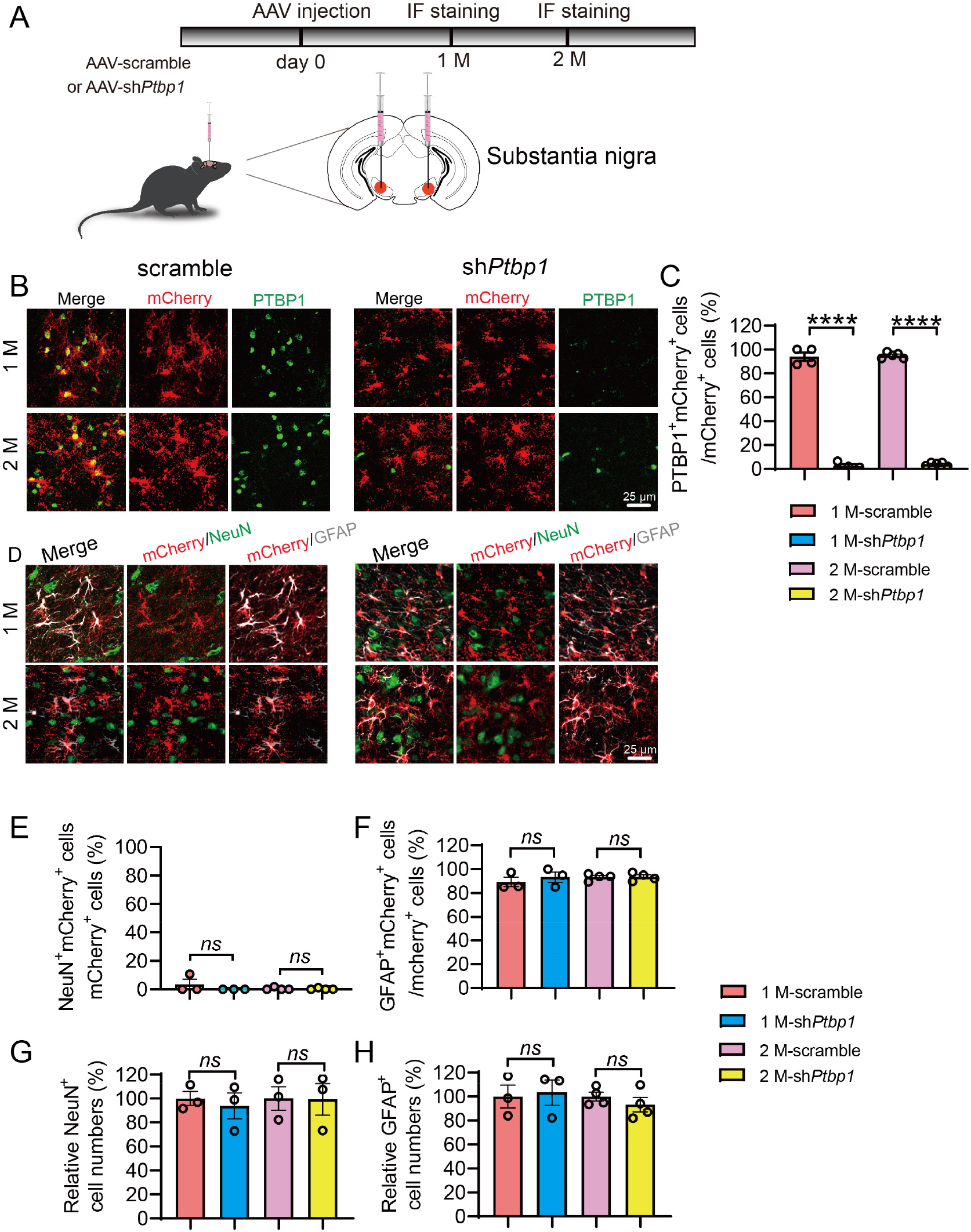
Downregulation of PTBP1 cannot convert astrocytes into neurons in the mouse substantia nigra. (**A**) Schematic of study design. (**B** and **C**) Confocal analysis of PTBP1 expression in mCherry^+^ cells. (B) Representative images, (C) quantification. n = 4 to 5 animals, one-way ANOVA with Tukey’s multiple comparisons. (**D** to **H**) Confocal analysis of fluorescent cells. (D) Representative images, (E-H) quantification. n = 3 to 4 animals, one-way ANOVA with Tukey’s multiple comparisons. All quantified data are represented as mean ± SEM; ****p <0.0001; *ns*, not significant. Figure 7—source data 1 to 5, source data for Figure 7C, 7E, 7F, 7G, 7H.

## Discussion

Having efficiently downregulated PTBP1 in the astrocytes in combination with cell lineage and transcriptomic analyses, we fail to observe any glia-to-neuron conversion induced by PTBP1 knockdown in multiple rodent brain regions at different ages, under either physiological or pathological conditions associated with AD. Similar to what we have found, a recent study also failed to replicate the reported astrocyte-to-neuron conversion as a result of PTBP1 knockdown (Wang et al., 2021). In the same study, Wang et al observed a dramatically increased number of NeuN^+^mCherry^+^ cells in the mouse brain transduced with astrocyte-restrictive AAV expressing mCherry reporter and NeuroD1. They therefore employed stringent lineage tracing strategies to investigate the origin of the increased NeuN^+^mCherry^+^ cells, and revealed that they are in fact endogenous neurons that were experimentally labeled with mCherry due to altered cell type specificity of the AAV virus induced by NeuroD1 overexpression. Because we found no evidence for neuronal generation induced by PTBP1 downregulation, it is therefore unnecessary to perform the lineage tracing analysis.

The discrepancies between the findings from Fu-Yang labs and our and Zhang labs are not likely due to differentially viral toxicities, because similar titers of AAV were used in all studies; instead, the differential leakage of astrocytic labeling systems would be more likely. Our AAV driven by the long form *hGFAP* promoter showed extremely low leaky expression in neurons. Apparent neuronal leakage of Cre expression in neuronal cells has been well documented in the *mGfap-Cre* line (Wang et al., 2021), which was utilized in Qian et al. study. Specifically, a control for PTBP1 downregulation was missed in the experiments evaluating astrocyte to neuron conversion *in vivo* (Qian et al., 2020), (Fig. 2b-h and Fig. 3). Although the information pertinent to the *GFAP* promoter in AAV-CasRx plasmids was completely missing in Zhou et al study, the information from Addgene (https://www.addgene.org/154001/ and https://www.addgene.org/154000/) shows that these plasmids comprise a short form *GFAP* promoter. Wang et al, however, was not able to downregulate PTBP1 and hence trigger glia-to-neuron conversion using the same AAV-CasRx system (Wang et al., 2021). In addition, we observed that PTBP1 predominantly localizes in nucleus, a subcellular compartment for RNA-binding proteins, whereas PTPB1 was detected all over the cell without a specific subcellular localization in Zhou et al. study (Zhou et al., 2020).

The definitive aim of neuronal regeneration is to improve, if not completely restore, brain functions deteriorated in neurodegeneration. Synaptic failure, a primary cause for cognitive impairment in AD, ought to be rescued if nascent neurons would be replenished from the resident glia cells. Our results demonstrate that PTBP1 downregulation cannot improve cognitive and synaptic function through any mechanisms including the neuronal regeneration. However, we do not exclude the possibility that PTBP1 downregulation could improve other brain functions such as motor function, which has been shown previously (Qian et al., 2020; Zhou et al., 2020). Future experiments are required to more carefully and in greater details test the strategy targeting PTBP1 in treating neurodegenerative diseases in not only rodents but also primates.

## Materials and methods

### Key Resource Table

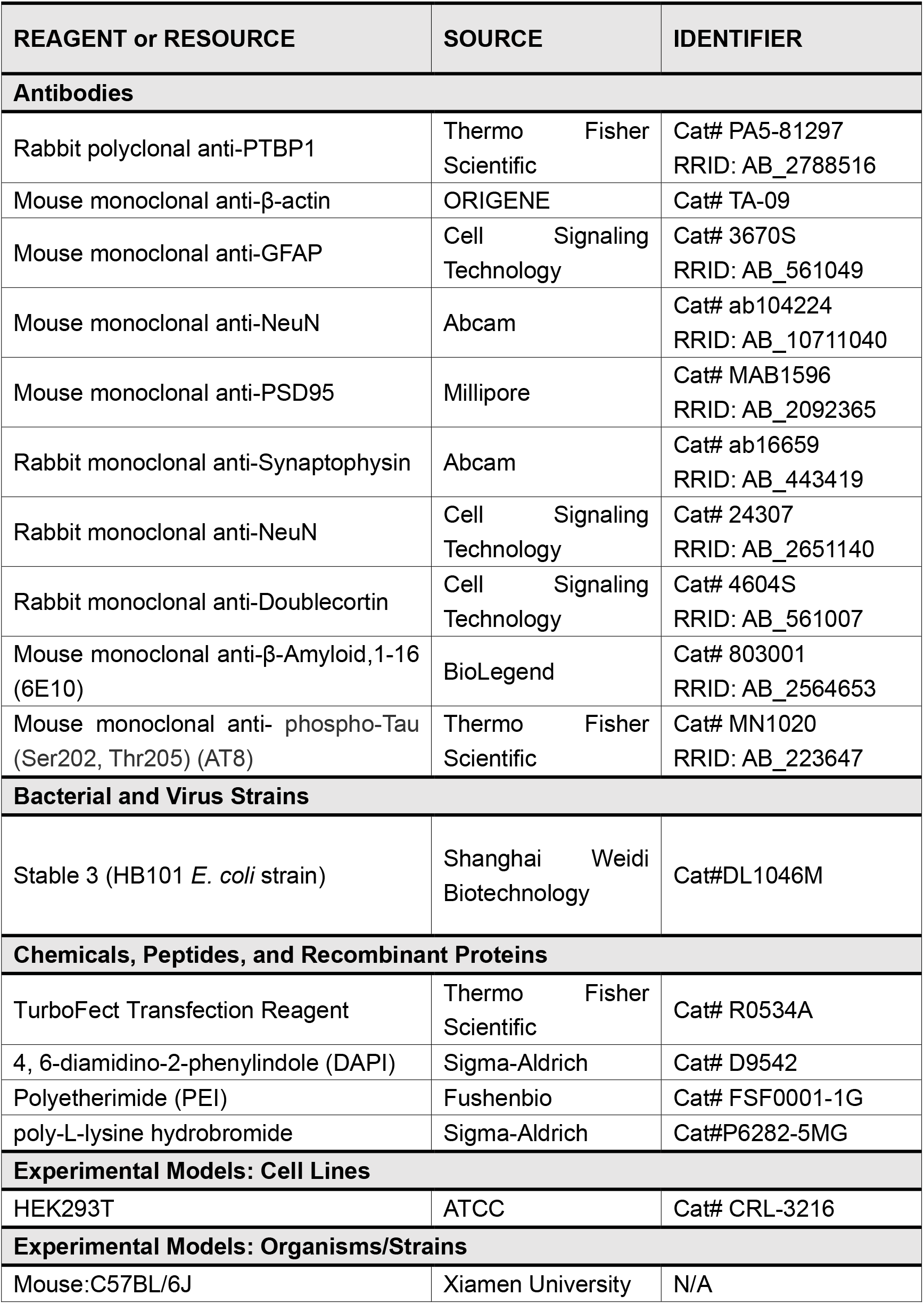
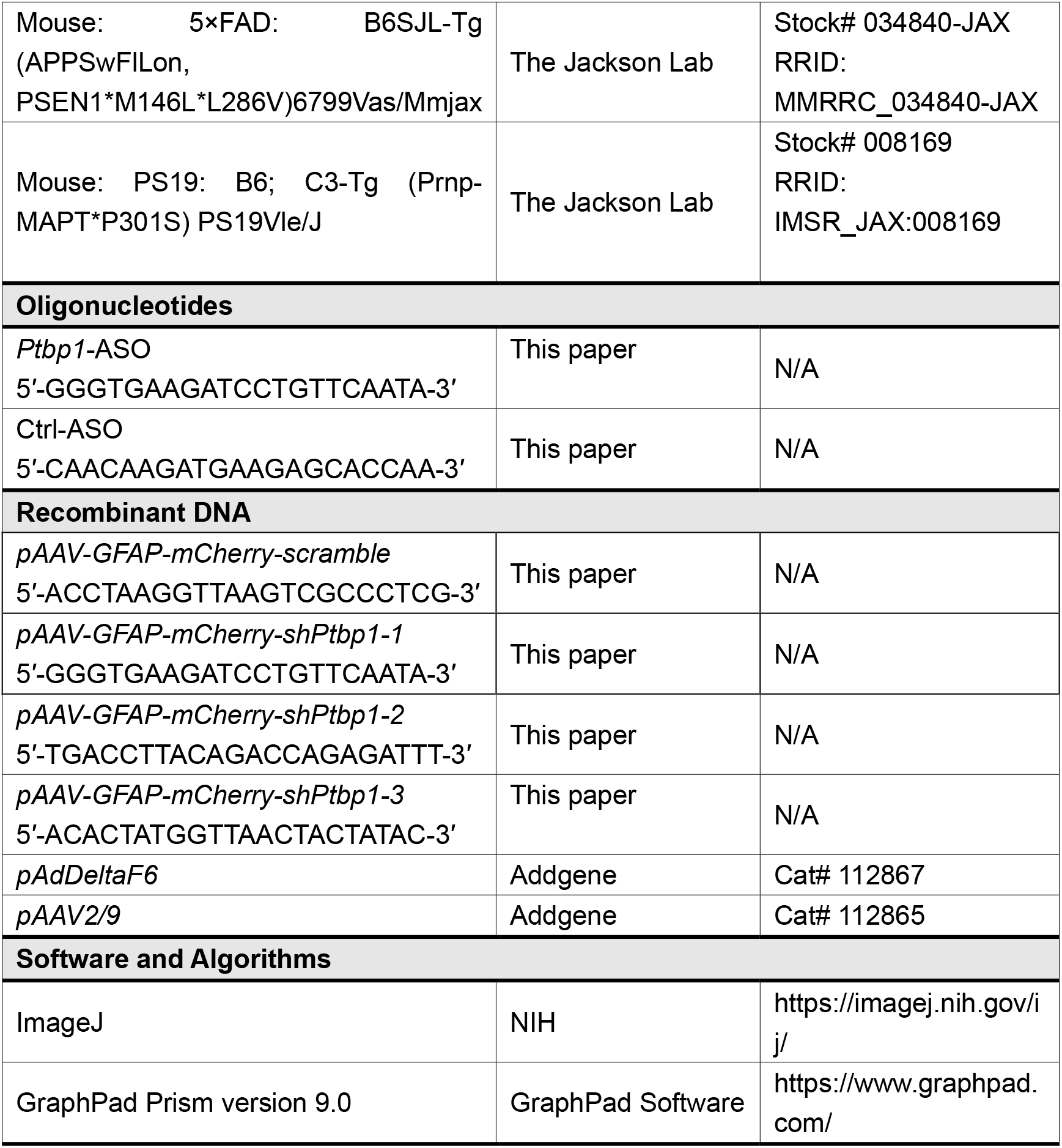

### Animals

Wild-type C57BL/6J mice were obtained from the Laboratory Animal Center at Xiamen University. 5×FAD mice (The Jackson Lab: Stock #034840-JAX); PS19 mice (The Jackson Lab: Stock #008169) were obtained from the Jackson Laboratory (Ellsworth, ME, USA) and were backcrossed into the C57BL/6J for 10 generations. Only male mice were used in behavioral tests, while both male and female mice were used in immunohistochemistry. Mice were randomly grouped by genotype and age. Experiments were conducted and analyzed in a double-blind manner. All animal studies were performed according to the protocols approved by the Institutional Animal Care and Use Committee of Xiamen University.

### Isolation and Culture of Primary Mouse Astrocytes

A modified protocol was performed to use (Schildge et al., 2013). Cortical and hippocampal tissues were dissected from P1-P2 pups and dissociated with 0.05% trypsin for 30 min at 37 °C. Tissues were centrifuged for 5 min at 500 g and mechanically dissociated in DMEM/F12 growth media conaining 20% FBS and 1% penicillin/streptomycin. Cells were passed through a 70 μm cell strainer, centrifuged at 500 g for 3 min, and resuspended in the growth media. Cells were plated into 0.1% poly-L-lysine (Sigma-Aldrich, #P6282) coated 175 cm^2^ flasks for proliferation. To examine the efficiency of AAV-mediated PTPB1 downregulation *in vitro,* cells were re-plated into 6-well plates, transduced with the AAV and lysed 7 days after the transduction.

### Western blot

Western blot was performed as previously described (Zeng et al., 2019). Briefly, primary cultured astrocytes were lysed in RIPA buffer (150 mM NaCl, 50 mM Tris-HCl [pH 8.0], 2 mM EDTA, 1% NP-40, 0.1% SDS, 0.5% sodium deoxycholate) containing protease inhibitor cocktail (Roche, #04693132001). Equal amounts of total proteins (20 μg per sample) from the lysates were resolved by SDS-PAGE. The samples were then probed with the primary antibodies against PTBP1(rabbit; 1:1000; Thermo Fisher Scientific; #PA581297) and β-actin (mouse; 1:5000; ORIGENE; #TA-09), followed by HRP conjugated secondary antibodies against rabbit or mouse IgG (1:5000; Thermo Fisher Scientific; #7074S or #7076S).

### Vectors and AAV production

To generate astrocyte specific AAV expression vectors, shRNAs targeting mouse *Ptbp1* or scrambled shRNA was inserted into an AAV vector driven by the 2.2 kb long human *GFAP* promoter *(pAAV-GFAP-mCherry-miR30E)* (VectorBuilder: https://www.vectorbuilder.cn/design/retrieve.html, VB200627-1109ntb). The sequences of shRNA-2 and −3 were obtained from Vectorbuilder (https://www.vectorbuilder.cn/design/pRP_shRNA.html), while shRNA-1 sequence was designed based on a recent study (Qian et al., 2020). The target sequences of shRNAs and scramble were as follows: scramble: 5’-ACCTA AGGTT AAGTC GCCCT CG-3’; sh*Ptbp1*-1:5’-GGGTG AAGAT CCTGT TCAAT A-3’; shP*tbp1*-2: 5’-TGACC TTACA GACCA GAGAT TT-3’; *shPtbp1-3:* 5’-ACACT ATGGT TAACT ACTAT AC-3’.

AAV viruses were produced following a well-established protocol with minor modifications (Challis et al., 2019). Briefly, HEK293T cells (ATCC, #CRL-3216) were transfected with the pAAV-shRNA, pAAV2/9 Helper (Addgene, #112867), and pAd-deltaF6 (Addgene, #112865) plasmids. Three days post transfection, viral particles were collected from the media and lysates of the cells, and purified by iodixanol gradient ultracentrifugation in a Beckman L-100XP centrifuge with type 70 Ti rotor at 350,000g for 2 hours and 25 min at 18 °C.

### Stereotactic injection

Stereotactic injection was performed as described previously (Wang et al., 2013; Zhao et al., 2015). Briefly, after anesthetization with Avertin, the experimental mice were shaved to remove the head hairs and secured in an automated stereotaxic injection apparatus (RWD Life Science). Ctrl-ASO or *Ptbp1-* ASO (2 μl, 2.5 μg/μL) were injected into the following coordinates: for hippocampal den-tate gyrus. −2.5 mm anteroposterior (AP) from the bregma, ±2.0 mm mediolateral (ML), −2.4 mm dorsoventral (DL); The target sequences of ASO and Ctrl were as follows: PTB-ASO: 5’-GGGTG AAGAT CCTGT TCAAT A-3’; Ctrl-ASO: 5’-CAACA AGATG AAGAG CACCA A-3’; AAV-*shPtbp1* or AAV-scramble (2 μl, titer 1×10^12^ viral genomes/mL) viruses were injected into the following coordinates: for hippocampal dentate gyrus, −2.5 mm anteroposterior (AP) from the bregma, ±2.0 mm mediolateral (ML), −2.4 mm dorsoventral (DL); for striatum, −0.8 mm AP from the bregma, ±1.6 mm ML, −2.8 mm DV; for substantia nigra, −3.0 mm AP from the bregma, ±1.2 mm ML, −4.6 mm DL.

### Immunofluorescence

Mice were anesthetized with Avertin followed by intracardial perfusion with PBS and 4% paraformaldehyde (PFA) in PBS. Mouse brains were dissected quickly, post-fixed with 4% PFA overnight at 4 °C, and dehydrated with 30% sucrose in PBS until it completely settled. The brains were then encased in optimal cutting temperature compound (Sakura; #4583), and cut into serial frozen sections (30-40 μm) using a cryostat (Leica; #CM1950). The sections were permeabilized and blocked with PBS containing 0.3% Triton X-100 and 10% normal donkey serum for 45 min at room temperature. The sections were incubated with the following primary antibodies at 4 °C overnight: mouse anti-GFAP (1:500; CST; #3670S), rabbit anti-NeuN (1:500; CST; #24307), mouse anti-NeuN (1:1000; Abcam; #ab104224), rabbit anti-PTBP1 (1:200; Thermo Fisher Scientific; #PA5-81297), rabbit anti-Doublecortin (1:500; CST; #4604S), moues-anti-PSD95 (1:100; Millipore; #MAB1596), rabbit anti-Synaptophysin (1:200; Abcam; AB_443419), mouse anti-Aβ (6E10) (1:500; BioLegend; #803001) and mouse anti-Phospho-Tau (Ser202, Thr205) (AT8) (1:200; Thermo Fisher Scientific; #MN1020). Donkey derived secondary antibodies conjugated with Alexa Fluor-488 or −647 (1:500; Thermo Fisher Scientific; #A-11034) (1:500; Thermo Fisher Scientific; #A-31571) were used for fluorescence, and DAPI (1μg/ml, Sigma-Aldrich, #D9542) was used to counterstain nuclei. Images were captured using a Leica SP8 or Zeiss LSM880 confocal microscope and subjected to quantification with ImageJ software [National Institutes of Health (NIH), https://imagej.nih.gov/ij/]

### Phospho-Tau examination

The levels of phosphor-Tau (T181) were determined by Single Molecule Immune Detection method (Astrobio).

### Behavioral tests

#### Morris water maze (MWM)

MWM tests were performed in a 1.2 m diameter circular tank filled with opaque water at 22 °C, using a modified protocol (Du et al., 2018). The walls surrounding the tank were marked with bright, contrasting shapes which serve as spatial reference cues. A fixed platform (10 cm diameter) was placed in a selected target quadrant. During training, the platform was submerged and the mice were placed into the maze at one of four points randomly facing the wall of the tank. Mice were allowed to search for a platform for 1 min; if the mice were unable to find the platform, they were gently guided to the platform for 10 s. Two trials a day were conducted with a minimum of a 1 hour intertrial interval. On day 6, the hidden platform was removed and probe test was performed. Escape latency to find the platform during training and time spent in target quadrant during probe test was recorded and analyzed by CleverSys TopScanLite (Clever Sys, Reston, VA, USA).

#### Novel object recognition (NOR)

NOR consists of three phases: habituation, training and test (Zhao et al., 2019). On day 1, mice were habituated to an open field box (40cm x 40cm x 40cm) for 5 min. On day 2, two same objects were placed in two diagonal corners of the box, and mice were put into the box and explored for 8 min. 24 hours later (day 3), one of the object was replaced with a novel object, and then mice were put back to the box and allowed to explore for 8 min. Cumulative time of each mouse spent on exploring each object was recorded by TopScanLite (Clever Sys, Reston, VA, USA). The discrimination index was calculated as the following: discrimination index = novel object exploration time/ (novel object exploration time + familiar object exploration time).

#### Contextual fear conditioning (CFC) test

CFC tests were performed based on a modified protocol (Shoji et al., 2014). On day 1, mice were placed into a conditioning box with a metal plate, and allowed to explore the box freely for 2 min as habitation. Thereafter, mice were received a 2 s foot electric shock (0.5 mA) for three times, with a 1 min interval between each time. Mice were left in the box for an additional min after the final electric shock. For the contextual test, mice were re-exposed to the same conditioning box for 5 min, 14 days after training. Time of freezing behavior for each mouse was analyzed by CleverSys FreezeScan (Clever Sys, Reston, VA, USA). Freezing % was calculated as the following: freezing %= (freezing time / total time) during the test – (freezing time / total time) during habitation.

#### Electrophysiology

For analysis of long-term potentiation (LTP), ex-vivo hippocampal slices were prepared from 10-month-old wild type (WT) and 5×FAD mice at three months post the transduction of AAV-sh*Ptbp1* or AAV-scramble. All the subsequent procedures are same as described previously (Zheng et al., 2021).

## Acknowledgements

We thank Baoying Xie, Haiping Zheng, Yixian Gao, Xiang You, Qingfeng Liu and Jingru Huang at Xiamen University for technical assistance.

## Author contributions

TG, HX and YZ conceived and designed the study, TG and XP performed behavioral analysis, XP performed and analyzed electrophysiology experiments, TG, XP, GJ, DZ carried out immunofluorescence staining and confocal analysis, JQ, LS and ZW provided helpful discussion, TG, HX and YZ wrote the manuscript. All authors approved the manuscript.

## Data and materials availability

Quantification results are displayed as mean ± SEM. Samples sizes are comparable to those described previously in similar studies (Qian et al., 2020; Wang et al., 2021), although statistical methods were not used to predetermine sample sizes in this study. Details for sample sizes and statistical tests can be found in the figure legends. Statistical analyses were performed with GraphPad Prism software (version 9.0, https://www.graphpad.com/). Differences were assessed by unpaired t tests or one-way ANOVA where appropriate. P values < 0.05 were considered as statistically significant.

