## Supplementary figures for "Downregulating PTBP1 fails to convert astrocytes into hippocampal neurons and to alleviate symptoms in Alzheimer’s mouse models"

### Supplementary Materials

**Figure 1—figure supplement 1. *Ptbp1*-ASO does not specifically enter astrocytes and fails to promote neuronal generation in the mouse hippocampus**

**Figure 2—figure supplement 1. AAV-medicated PTBP1 downregulation fails to convert hippocampal astrocytes into neurons in 5×FAD mice 1 month after the AAV transduction**

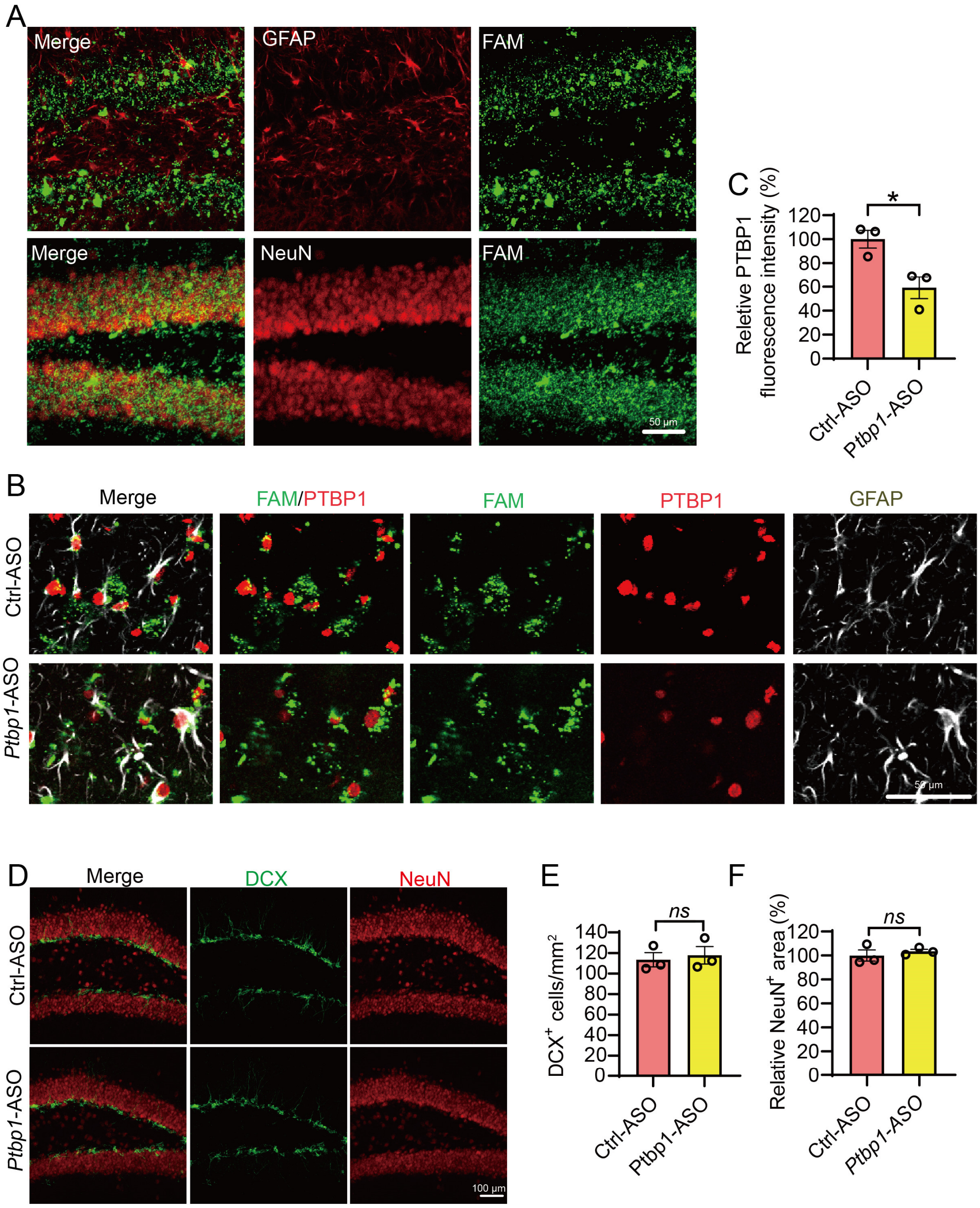


**Figure 1—figure supplement 1. *Ptbp1*-ASO does not specifically enter astrocytes and fails to promote neuronal generation in the mouse hippocampus**

(**A**) Confocal analysis of cellular distribution of FAM-ASO in the hippocampus. (**B** and **C**) Confocal analysis of PTBP1 expression in $FAM^+GFAP^+$ cells. (B) Representative images, (C) quantification. n = 3 animals per group, unpaired t-test. (**D** to **F**) Confocal analysis of $DCX^+$ cells and $NeuN^+$ cells. (D) Representative images, (E and F) quantification. n = 3 animals per group, unpaired t-test. All quantified data are represented as mean ± SEM; $^*p < 0.05$; *ns*, not significant. Figure 1 figure supplement 1—source data 1 to 3, source data for Figure 1—figure supplement 1C, 1E, 1F.

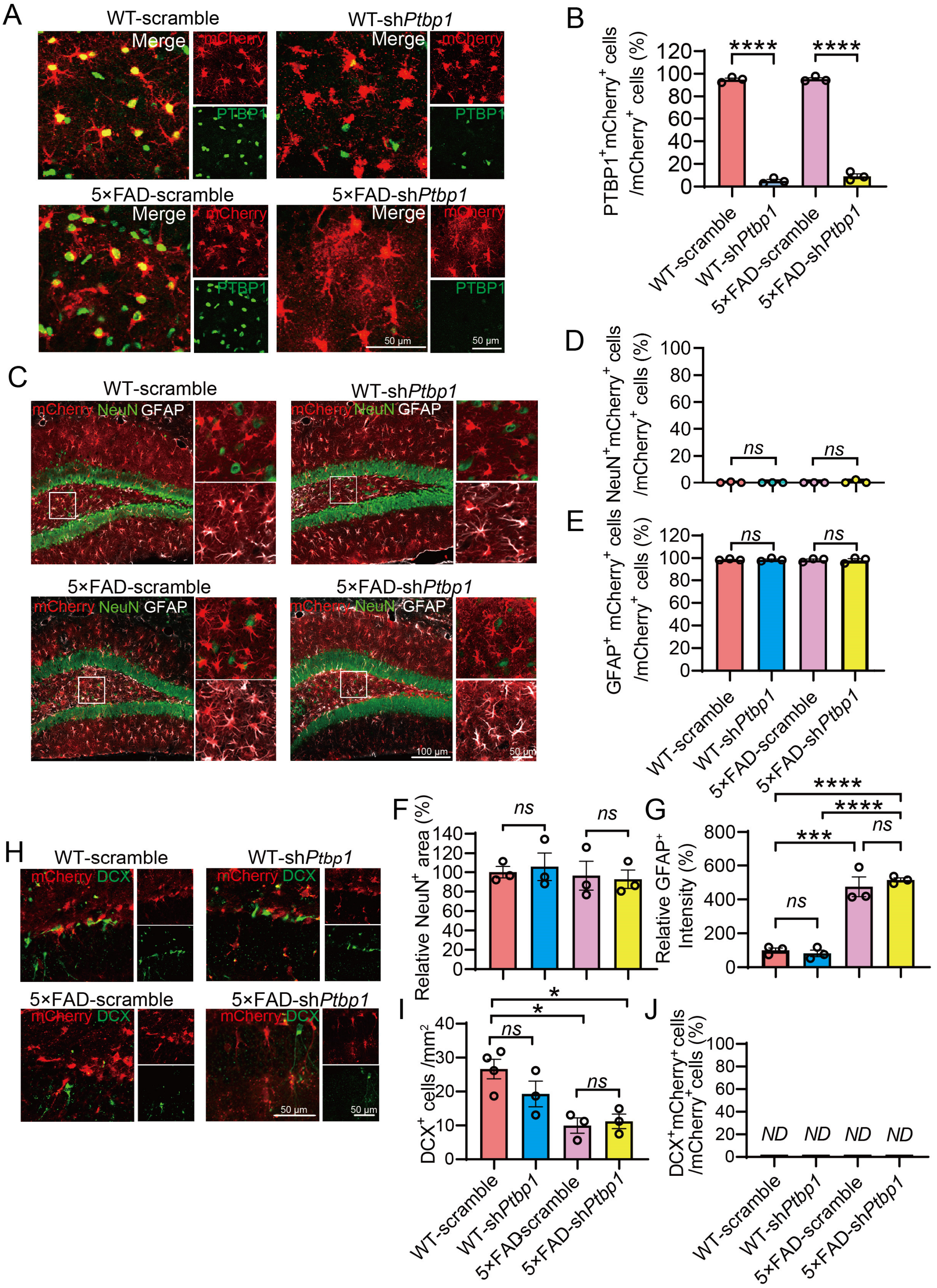


**Figure 2—figure supplement 1. AAV-medicated PTBP1 downregulation fails to convert hippocampal astrocytes into neurons in 5×FAD mice 1 month after the AAV transduction**

(**A** and **B**) Confocal analysis of PTBP1 expression in mCherry+ cells. (A) Representative images, (B) quantification. n = 3 animals per group, unpaired t-test. (**C** to **G**) Confocal analysis of fluorescent cells. (C) Representative images, (D-G) quantification. n = 3 animals per group, one-way ANOVA with Tukey's multiple comparisons. (**H** to **J**) Confocal analysis of $mCherry^+$ cells and $DCX^+$ cells. (H) Representative images (I and J) quantification. n = 3 to 4 animals, one-way ANOVA with Tukey's multiple comparisons. All quantified data are represented as mean ± SEM; *p <0.05; ***p <0.001; ****p <0.0001; *ns*, not significant; ND, undetectable. Figure 2-figure supplement 1—source data 1 to 7, source data for Figure 2—figure supplement 1B, 1D, 1E, 1F, 1G, 1I, 1J.
